# Tropical Race 4 and Race 1 strains causing Fusarium wilt of banana infect and survive in *Heliconia* species and ornamental bananas

**DOI:** 10.1101/2023.12.11.571145

**Authors:** Einar Martínez de la Parte, Harold J.G. Meijer, Mauricio Guzmán Quesada, Claudiana Carr Rodriguez, Silvia Masis Jimenez, Luis Pérez Vicente, Gert H.J. Kema

**Affiliations:** Laboratory of Phytopathology, Wageningen University & Research, The Netherlands; Instituto de Investigaciones de Sanidad Vegetal (INISAV), Cuba; BU Biointeractions & Plant Health, Wageningen University & Research, The Netherlands; Cropland Bioscience, Costa Rica; Departamento de Fitoprotección, Corporación Bananera Nacional (CORBANA S.A.), Limón, Costa Rica; Laboratorio de Cultivo de tejidos, Coordinación de Fisiología, Clima y Producción, Corporación Bananera Nacional (CORBANA S.A.), Limón, Costa Rica

**Keywords:** *Heliconia*, *Fusarium odoratissimum*, TR4, abacá, secondary hosts

## Abstract

Fusarium wilt of banana (FWB), caused by soilborne *Fusarium* spp., is a major global threat to the cultivation of bananas. In addition to persistent chlamydospores, weeds are a reservoir of the causal agents. However, it remains unclear whether other Zingiberales species, which are grown in the same geographic regions, also can serve as hosts for *Fusarium* spp. that cause FWB. Greenhouse assays were conducted to investigate whether *Fusarium phialophorum* (Race 1; pathogenic to Gros Michel banana) and *Fusarium odoratissimum* (TR4; pathogenic to Cavendish banana) can infect three *Heliconia* species, two ornamental banana species or *Musa textilis* (abacá). *Heliconia latispatha, Musa balbisiana*, and *Musa coccinea* displayed external symptoms after inoculation with TR4, while inoculation with Race 1 caused symptoms in *H. latispatha, H. psittacorum, M. coccinea*, and *M. velutina*. Isolates recovered from distinct organs of all studied plant species were characterized and re-isolated strains caused FWB symptoms in Gros Michel and Cavendish banana plants, and their rhizome discolored area scores were similar to the reference strains. The susceptibility of some ornamental species and the presence of *Fusarium* strains as asymptomatic endophytes in others, with remaining pathogenicity, call for a revision of the race nomenclature and the current containment protocols for FWB.

## INTRODUCTION

The genus *Musa* belongs to the Musaceae family of the order Zingiberales and comprises many wild and seeded banana species as well as all seedless and therefore edible varieties. Cultivated bananas are a major staple food for millions of people in many countries and dominate the global fruit commerce (FAO, 2022; Ploetz, 2015). The genus *Musa* is divided into two sections, *Callimusa* and *Musa*, with the latter including most edible banana cultivars (Häkkinen, 2013), which are diploid or triploid hybrids of *Musa acuminata* (AA, 2n=22) or from hybridization with *Musa balbisiana* (BB, 2n=22) (Ploetz, 2006). Only a minor group of cultivars, which included Fe’i bananas, is derived from species of the *Callimusa* section (Häkkinen, 2013). This section also includes *M. textilis* (also known as abacá) which is appreciated as source of cordage fiber. Ornamental banana species can be found in both of the aforementioned sections and typically have relatively few fruits and are best known for their brightly colored bracts (Häkkinen, 2013; Häkkinen and Väre, 2008). Another group of ornamental species, *Heliconia* spp. also belong to the Zingiberales, but to the family of the Heliconeaceae. They are very popular neotropical ornamentals characterized by colorful inverted flowers (Gómez-Merino et al., 2018).

Fusarium wilt of banana (FWB) is a devastating vascular disease that severely impacts international banana production (Staver et al., 2020) and jeopardizes food security and livelihoods in regions that rely on banana cultivation (Fones et al., 2020; Steinberg and Gurr, 2020). The disease is caused by a diverse array of *Fusarium* species (Maryani et al., 2019; Mostert et al., 2022; Ordóñez et al., 2015; van Westerhoven et al., 2023). They invade the vascular system of banana plants after root infection, obstructing water and nutrient transport and ultimately leading to wilting, chlorosis and plant death (Pegg et al., 2019). To date, three different races of the FWB pathogen have been described based on the pathogenicity to reference banana cultivars, named Race 1, Race 2 and Race 4, with the latter subdivided into Tropical Race 4 (TR4) and Subtropical Race 4 (Pérez-Vicente, 2004; Ploetz, 2015). A pathogen population causing wilt in *Heliconia* spp., was described as Race 3 (Waite, 1963), but it is no longer considered part of the *Fusarium* complex infecting banana (Ploetz, 2015). The FWB pathogens also reside and survive in non-banana plant species (Waite and Dunlap, 1953; Pittaway, Nasir, and Pegg 1999; Hennessy et al., 2005; Martínez de la Parte, 2023) which contributes to their long-term survival in the field. Some *M. velutina* and *M. textilis* cultivars have been included in germplasm screenings for resistance to TR4 (García-Bastidas, 2019; Li et al., 2015; Zuo et al., 2018), but others, such as *M. coccinea*, have never tested. Additionally, *Fusarium* isolates infecting *M. textilis* have not been used in disease evaluations of bananas (Borines et al., 2007; Dyah, Purwati and Hidayah 2008). Thus, the understanding of the potential role of *Heliconia* spp. and the aforementioned non-edible *Musa* spp., in addition to nonrelated weed species (Martínez de la Parte et al., 2023), in the epidemiology and survival of FWB pathogens remains limited. Since these Zingiberales species are grown in the same geographical areas as bananas it is crucial to understand their possible role in the FWB epidemiology to support improved disease management strategies.

## MATERIALS AND METHODS

### Plant material

Tissue culture plants of *Heliconia latispatha* Benth., *Heliconia psittacorum* L., *Heliconia rostrata* Ruiz & Pav., *Musa coccinea* Andrews, *M. velutina* H. Wendl. & Drude, *M. textilis* Nee, and *cv*. Gros Michel (AAA) were propagated at the tissue culture laboratory of the Corporación Bananera Nacional (CORBANA S.A.), Costa Rica. Tissue culture banana plants *cv*. Grand Naine (AAA) were obtained from Vitropic (Saint Mathieu-de-Tréviers, France). Upon arrival at Wageningen University & Research (WUR, The Netherlands), the plants were transferred from transport plastic boxes to 1L pots containing a standard soil (Swedish sphagnum peat 20%, Baltic peat 30%, garden peat 30%, beam structure 20%, grinding clay granules 40.6 Kg/m3, Lime + MgO 2.5 Kg/m3, PG-Mix-15-10-20 0.8 Kg/m3) from the WUR-Unifarm greenhouse facility. The potted plants were then acclimatized under plastic to maintain high humidity conditions for two weeks in an environmentally controlled greenhouse compartment and thereafter grown for ∼2.5 months prior to inoculation (28±2°C, 16h light, and ∼85% relativity humidity). During the experiment, plants were watered daily and fertilized three times per week (NH_4_^+^-1.2 mM/L, K^+^-7.2 mM/L, Ca^2+^-4 mM/L, Mg^2+^-1.82 mM/L, NO_3_^-^-12.4 mM/L, SO_4_^2-^-3.32 mM/L, H_2_PO_4-_-1.1 mM/L, Mn^2+^-10 µMol/L, Zn^2+^-5 µMol/L, B-30 µMol/L, Cu^2+^-0.75 µMol/L, Mo-0.5 µMol/L, Fe/DTPA-50/3%, Fe-EDDHSA-50/3%, pH=5.8).

### Inoculum production and inoculation methods

To produce inoculum conidial suspensions of the reference strains for TR4 and Race 1, *Fusarium odoratissimum* strain II5 and *F. phialophorum* strain CR1.1A (van Westerhoven et al., 2023), respectively, were produced in flasks containing 100 mL of mung beans broth (2 gr mung beans per 500 ml water) according to García-Bastidas et al., (2019). The flasks were incubated at 25°C, 150 rpm for five days, after which the concentration was adjusted to 1 × 10^6^ conidia.ml^−1^ prior to inoculation.

Inoculations were performed by wounding the roots in the soil using a soil scoop at two opposite sides of the plant and subsequent drenching with a conidia suspension of 200 ml of 10^6^ conidia/L per pot. For negative controls, we drenched each pot with 200 ml of water after damaging the roots, and inoculated Grand Naine and Gros Michel plants were used as positive controls for TR4 and Race 1, respectively. During the experiment six plants of each species were inoculated with Race 1 and nine with TR4, while three to five plants were used as negative control (Table S1). The experiment was repeated twice with a Randomized Complete Block Design.

### Disease progress and diagnosis of reisolated strains from plant inoculated with Race 1 or TR4

Plants were inspected weekly for any noticeable FWB symptoms compared to the non-inoculated treatment and the positive controls. Five plants per species were randomly selected 12 weeks post-inoculation, and three pieces (∼1 cm^2^) of plant material were collected from the roots, rhizome or corm and pseudostem of each plant. Sampled plant parts were extensively washed with demineralized water and then surface sterilized with 70% ethanol for five minutes in a laminar flow cabinet, then three times rinsed with sterile demineralized water and placed on Komada’
ss semi-specific media for *Fusarium* spp. (Komada, 1975). After six days, small fragments of the edges of fungal colonies with *Fusarium* morphology were collected and transferred to fresh potato dextrose agar (PDA) plates.

For diagnosis, fragments of the mycelium on PDA plates (five to seven days old) were collected with a sterile toothpick and transferred into a tube containing 20 µL of dilution buffer included in the direct Thermo Scientific Phire Plant Direct PCR Master Mix PCR master mix kit, which does not require separate DNA extraction (Thermo Scientific, Landsmeer, the Netherlands). The tube was vortexed for 10 s and incubated at 95°C for 5 min. One µL of the solution without mycelia was used as template in 20-µL PCR, which contains 10 µL of the Master Mix (2X) and 0.5 µL of each primer (10mM). The PCR program comprised an initial denaturation at 98°C for 5 min, followed by 35 or 40 cycles of denaturation at 98°C for 5 s, annealing at 62°C or 55°C (Table 1) for 5 s and extension at 72°C for 20 s, and a final extension at 72°C for 1 min. We used the primers of Dita et al., (2010), and Carvalhais et al., (2019) (Table 1) for final molecular diagnosis. The resulting amplicons were visualized after electrophoresis with a 100-bp ladder as a reference on 1.5% agarose gel by staining with ethidium bromide, and gels were photographed using the ChemidocTM MP image system (Bio-Rad Laboratories, Veenendaal, The Netherlands).

**Table 1.**
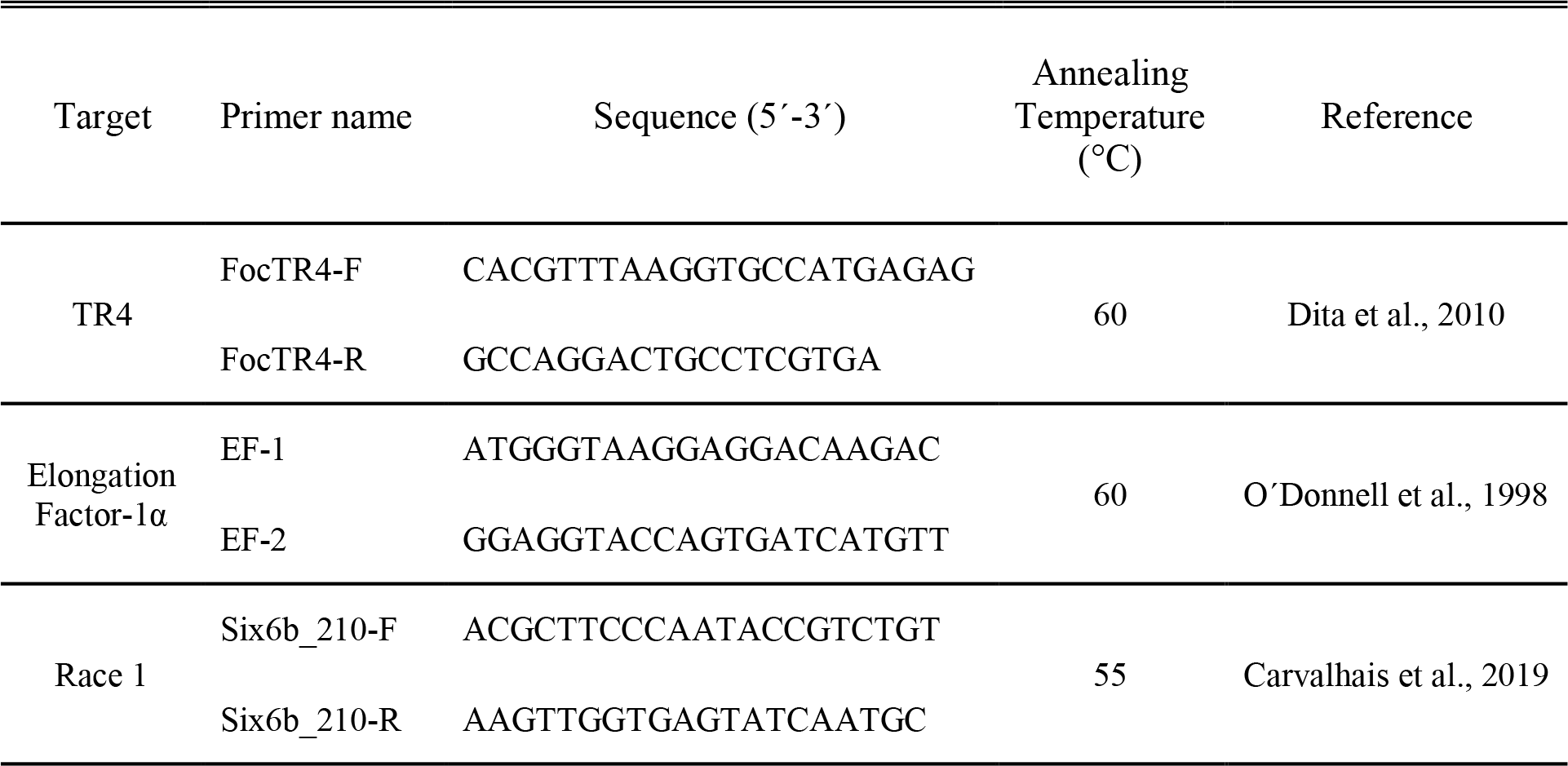
Primers for molecular diagnosis of re-isolated *Fusarium* strains.

### Pathogenicity of reisolated *Fusarium* isolates

From each plant species, we selected one isolate of Race 1 and one of TR4 that tested positive with the abovementioned molecular diagnostics and reisolated from the uppermost colonized part of the plant, for inoculations of banana following the aforementioned protocols. Twelve-week-old plants of Gros Michel and Grand Naine were inoculated with Race 1 and TR4 isolates, respectively, using the reference strains CR1.1A (Race 1) or II5 (TR4) as positive controls, and water-treated plants as negative controls (mock). For each re-isolate we inoculated in total seven Gros Michel plants and seven Grand Naine, in two separate experiments using a Randomized Complete Block Design. The rhizomes of the inoculated plants were cut transversally 11 weeks after inoculation, photographed and the Rhizome Discoloured Area (RDA) was calculated using ImageJ 1.52r software (National Institutes of Health, Bethesda, MD, USA). Individual RDA values were plotted using the web-tool BoxPlotR (Spitzer et al., 2014). The Kruskal-Wallis test was used to compare RDA values on Grand Naine and Gros Michel plants caused by the various isolates and Dunn’s multiple comparisons test was applied for multiple comparisons of variables at a P value of <0.05.

## RESULTS

To investigate the host range of the FWB pathogen among *Heliconia* spp. and the level of susceptibility of non-edible bananas used as ornamental or fiber crop, we conducted greenhouse tests where *H. latispatha, H. psittacorum, H. rostrata, M. coccinea, M. velutina* and *M. textilis* were challenged with inoculum of two reference isolates for Race 1 and TR4. Throughout all the individual phenotyping experiments conducted in this study, negative water controls - as well as the incompatible interaction between Grand Naine and Race 1 - never showed any external or internal FWB symptoms. The positive controls, Grand Naine vs. TR4 and Gros Michel vs. Race 1, developed severe FWB symptoms as expected (Figure 1).

**Figure 1.**
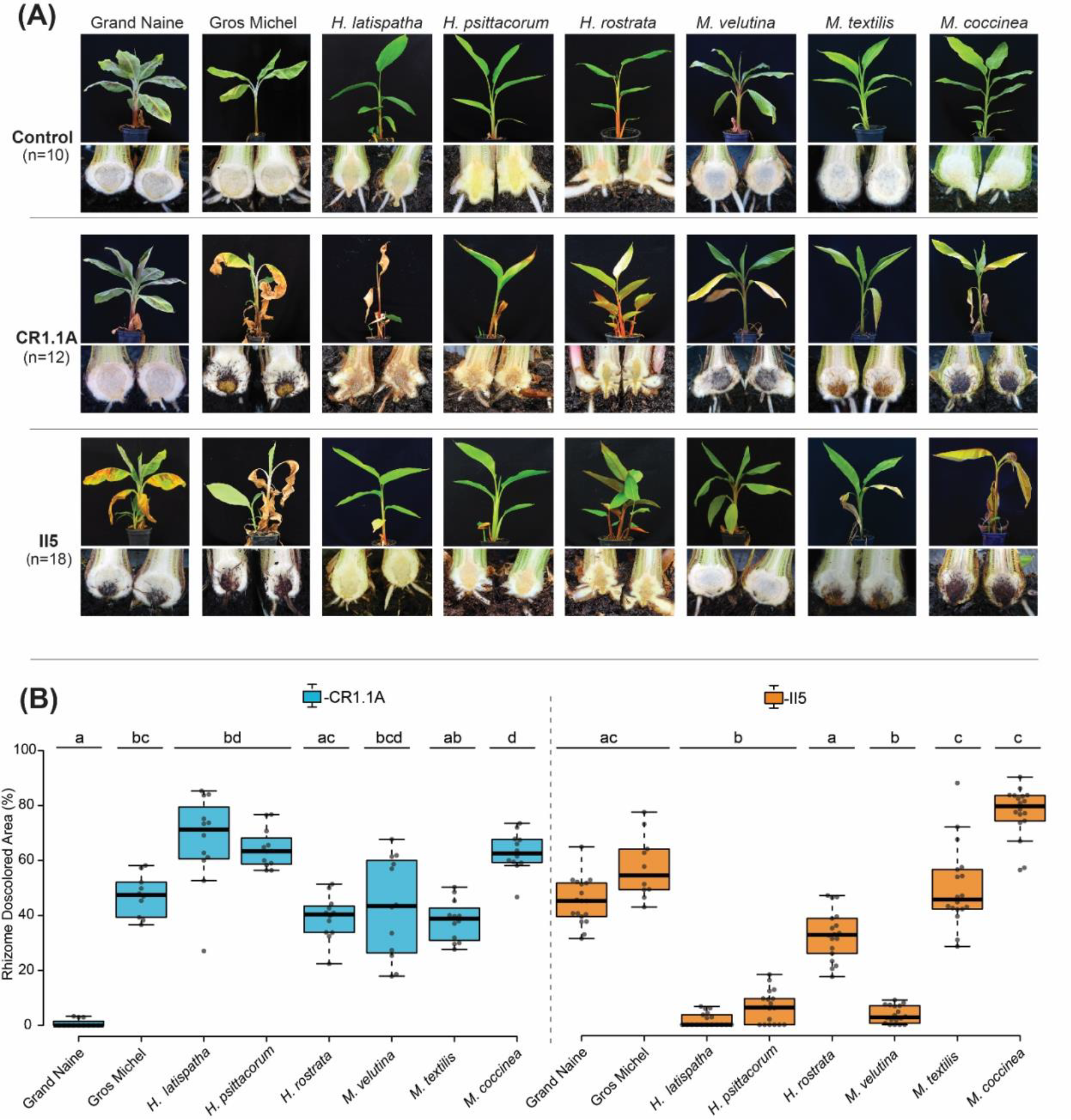
Differential responses of three *Heliconia* spp. and three non-edible *Musa* spp. after inoculations with *F. phialophorum* isolate CR1.1A representing Race 1 or *F. odoratissimum* isolate II5 representing TR4. A. External symptoms (upper rows) and cross sections of rhizomes (bottom rows) from representative plants 12 weeks after inoculation with mock (upper panel), Race 1 (middle panel) or TR4 (lower panel). B. RDA scores of inoculated plants. Statistically significant differences (p < 0.05) between median RDA values are indicated with different letters, according to Dunn’s multiple comparison test.

All plants of non-edible *Musa* spp. showed FWB symptoms typical for susceptible responses at 12 weeks after inoculation with Race 1 (Figure 1) with an average RDA value higher than 38% with *M. coccinea* being the most susceptible (RDA =62%). Both *M. coccinea* and *M. textilis* were also susceptible to TR4 with average RDA values of 77.8% and 53.2%, respectively. However, 35% of the *M. velutina* plants did not show any external or internal symptoms while the remainder of the plants showed the lowest scores among the tested *Musa* spp. (Figure 1) with an average RDA value of 3.8% at 12 weeks after inoculation.

Inoculation of the three *Heliconia* spp. with Race 1 caused wilting symptoms in *H. latispatha* and *H. psittacorum* but not in *H. rostrata*. The symptoms included leaf chlorosis, drying of leaves, weakened pseudostems that easily dislodged from the base upon pulling. However, all Race 1 inoculated *Heliconia* plants, including those of *H. rostrata*, showed necrosis in the roots and rhizomes (Figure 1). TR4 caused rhizome discoloration in all *Heliconia* species but never necrosis and only caused external symptoms in *H. latispatha*, similar to the symptoms caused by Race 1. Taken together, these results show that the *Heliconia* spp. and *M. velutina* plants are more susceptible to Race 1 than to TR4.

To further investigate the colonization of the tested *Heliconia* spp. and *Musa* spp. by Race 1 and TR4, various parts of inoculated plants were sampled to re-isolate the *Fusarium* strains. A total of 796 isolates was obtained (Table 2), re-isolates representative of the different plant organs per specie were confirmed as Race 1 or TR4 by diagnostic PCRs (Figure S1). The presence of Race 1 or TR4 strains was confirmed in the roots, rhizomes and pseudostems of *H. latispatha, M. coccinea* and *M. textilis*. We could not recover TR4 from the pseudostems of *H. psittacorum, H. rostrata* or *M. velutina*. Similarly, no Race 1 isolates could be recovered from the pseudostem of inoculated *H. rostrata* plants or from any tissue obtained from Grande Naine plants (Figure S1). Thus, we have demonstrated that causal agents of FWB are able to colonize the aboveground tissues of *H. latispatha, H. psittacorum, M. coccinea* and *M. textilis* plants.

**Table 2.** The number of recovered isolates from different plant parts of three *Heliconia* and *Musa* species after inoculation with Race 1 and TR4.

| Plant species | Race 1 |  |  | TR4 |  |  |
| --- | --- | --- | --- | --- | --- | --- |
|  | Roots | Rhizome | Pseudostem | Roots | Rhizome | Pseudostem |
| *Heliconia latispatha* | 25 | 24 | 15 | 24 | 8 | 5 |
| *Heliconia psittacorum* | 20 | 20 | 5 | 21 | 9 | - |
| *Heliconia rostrata* | 18 | 6 | - | 14 | 7 | - |
| *Musa coccinea* | 31 | 31 | 22 | 32 | 30 | 29 |
| *M. velutina* | 35 | 35 | 30 | 31 | 21 | - |
| *Musa textilis* | 31 | 32 | 17 | 34 | 28 | 10 |
| Gros Michel | 23 | 20 | 19 | 24 | 25 | 21 |
| Grand Naine | - | - | - | 34 | 32 | 30 |

To determine whether the pathogenicity of the recovered isolates was altered by passaging through these hosts, we used one confirmed Race 1 and TR4 re-isolate per plant species, to inoculate Gros Michel and Grand Naine banana plants. Isolates were selected from the uppermost colonized part and their pathogenicity did not significantly differ from the reference strains CR1.1A and II5, except the TR4 isolate recovered from *H. rostrata* which scored significantly lower than the reference strain *F. odoratissimum* II5 strain with an average RDA of 14.9%, (Figure 2).

**Figure 2.**
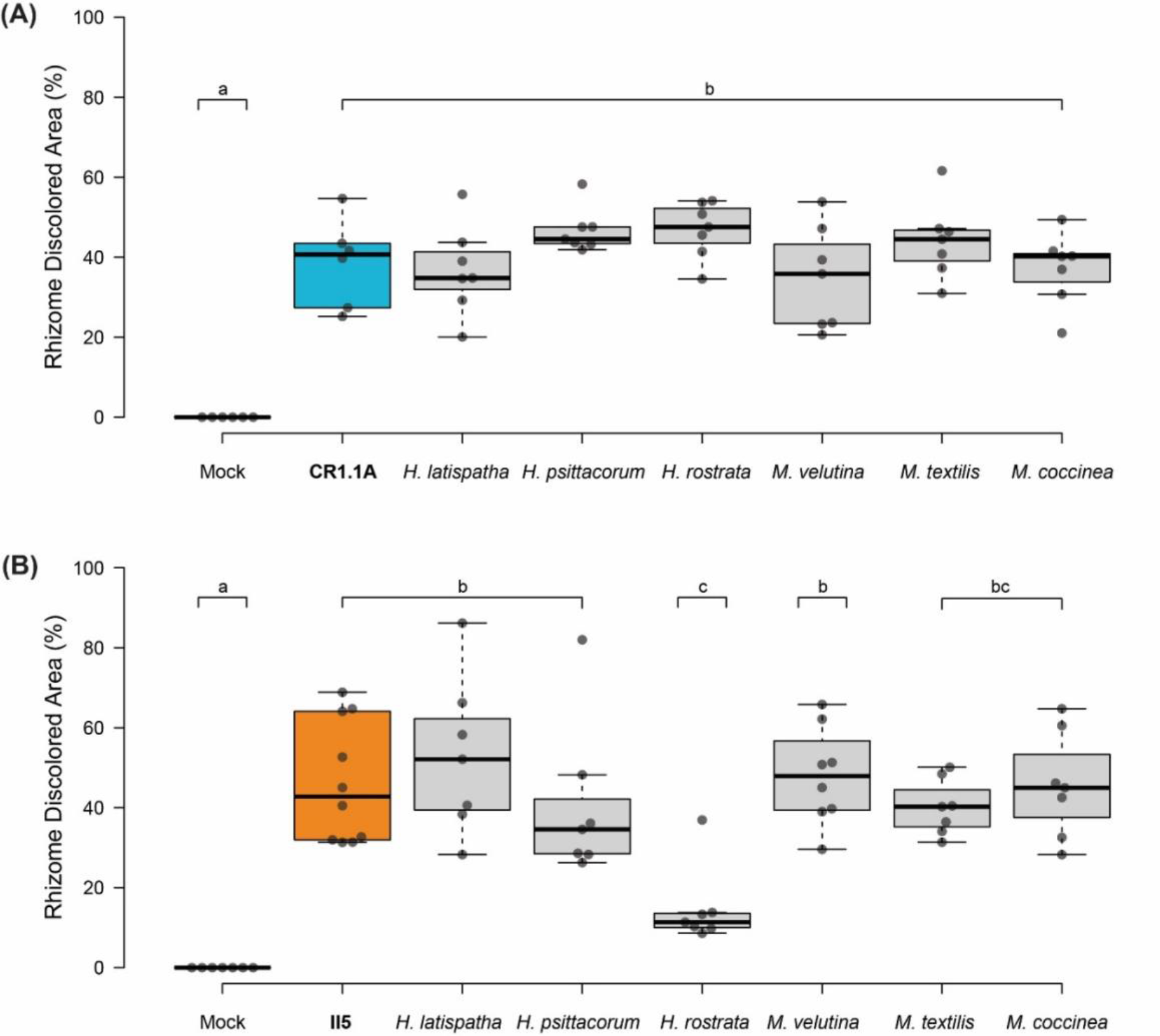
Phenotyping results of six re-isolated *Fusarium* strains from three *Heliconia* and three *Musa* spp. at 11 weeks post-inoculation. A. Race 1 isolates tested on Gros Michel and compared with the *F. phialophorum* Race 1 reference strain CR1.1A. B. TR4 isolates tested on Grand Naine and compared with the *F. odoratissimum* TR4 reference strain II5. Different letters indicate statistically significant differences (p < 0.05) between median RDA values, according to Dunn’s multiple comparison test.

## DISCUSSION

Various (tropical) plants and plant products are traded around the globe, often facilitating the unintended transport of associated pathogens or endophytes (Fones et al., 2020). The intensified movement and volume of traded plants and plant products over the last decades has resulted in an unprecedented spread of pathogens at a global scale, with the current dissemination of TR4 as one of the clearest examples (Drenth and Kema, 2021). This worrisome global spread, and particularly the recent cases in Latin America and Mozambique (Acuña et al., 2021; Garcia-Bastidas et al., 2020; Herrera et al., 2023; Westerhoven et al., 2022a), shows that FWB successfully disseminates despite implemented quarantine and prevention strategies (Westerhoven et al., 2022b). The recommended eradication and containment strategies, (Dita et al., 2018; Viljoen et al., 2020), largely ignore the presence of secondary hosts of the pathogen (Pegg et al., 2019). Thus, the identification of such alternative hosts could be useful in understanding pathogen persistence, inoculum accumulation, and subsequent spread to other areas (Hennessy et al., 2005; Martínez de la Parte et al., 2023).

*Heliconia* spp. are native to the tropical regions of the Americas, primarily Central and South America, with some species also found in the Caribbean and the Pacific islands (Gómez-Merino et al., 2018). In those regions, they are often present in rainforests, along riverbanks, and naturally growing alongside or near banana plantations. In addition, they are cultivated to produce cut flowers for the export in Latin-American and in some African countries (Linares-Gabriel et al., 2020), and are also used in landscaping parks and gardens (Gómez-Merino et al., 2018). Similar to bananas, Fusarium wilt is one of the most common diseases in *Heliconia* spp. (Castro et al., 2010). The causal agent of wilt in *Heliconia* spp. was initially described as Race 3 of the FWB pathogen (Waite, 1963) but is no longer considered as part of the causative *Fusarium* complex infecting banana (Ploetz, 2015). Recent studies evaluating the pathogenicity of *Fusarium* strains from *Heliconia* genotypes did not include testing on bananas (Castro et al., 2008; 2010). Nevertheless, only limited information is available on the potential infection of *Heliconia* by FWB-causing *Fusarium* spp. This is a significant knowledge gap when considering the potential impact of global dissemination of these species, particularly since it has been shown that weeds can contribute to FWB spread which requires altered disease management strategies (Hennessy et al., 2005; Martínez de la Parte et al., 2023; Pittaway et al., 1999; Su et al., 1986; Waite and Dunlap, 1953).

Our results are mostly consistent with, but significantly extend, early studies which reported that Race 1 strains cause symptoms on *H. psittacorum* (Rishbeth, 1957), *H. caribea* and *H. latispatha* (Waite, 1961), and that re-isolated strains were pathogenic on Gros Michel. Then, the re-isolated strains were obtained exclusively from the belowground plant organs (Rishbeth, 1957; Waite, 1961), and information on the genotypes of the studied isolates is lacking. Moreover, recent phylogenetic studies confirmed that Race 1 is polyphyletic and comprises a suite of *Fusarium* spp. (Maryani et al., 2019; van Westerhoven et al., 2023). Here, we demonstrated that passing Race 1 through *H. lathispatha* and *H. psittacorum* did not change the phenotype on Gros Michel. However, under our experimental conditions, *Heliconia* spp. exhibited higher susceptibility to Race 1 than to TR4. This could be attributed to the prolonged co-evolution of Race 1 with a plethora of *Heliconia* species in the Latin American and Caribbean region, which is considered the center of origin for these plants (Gómez-Merino et al., 2018; Malakar et al., 2022). We also showed for the first time that TR4 can colonize three *Heliconia* species, causing a range of symptoms. It is important to underscore that, in contrast to the other *Heliconia* spp., TR4 did not cause any external symptoms in *H. psittacorum* and *H. rostrata*, similar to the non-symptomatic colonization of various weeds and cover crops (Hennessy et al., 2005; Martinez de la Parte et al., 2023; Pittaway et al., 1999; Su et al., 1986; Waite and Dunlap, 1953), which urges adaptation of contingency and containment strategies upon the detection of TR4.

The *Musa* spp. we tested showed a varied reaction to the inoculation with the Race 1 and TR4 strains. The susceptibility to FWB of *M. textilis* was consistent with previous reports (García-Bastidas, 2019; Zuo et al., 2018), but we also observed that *M. coccinea* was equally susceptible to TR4 as to Race 1, and that *M. velutina* plants showed no wilting symptoms after inoculation with TR4, which confirms a previous report (Li et al., 2015). However, these authors neither examined the colonization of the different plant organs, nor phenotyped recovered strains. We show that passage of FWB-causing *Fusarium* spp. through these hosts generally does not affect pathogenicity to banana, except for strains recovered from *H. rostrata*, similar to the recently reported reduction of pathogenicity after passing TR4 through *Arachis pintoi* and *Portulaca oleracea* (Martínez de la Parte et al., 2023). The attenuated pathogenicity of TR4 after colonization of non-host plants and *H. rostrata* requires further studies, but these studies nevertheless underscore the need to consider the wider host range of TR4 in disease management strategies.

In conclusion, our results evidenced that non-edible *Musa* spp. and *Heliconia* spp., which are commonly used as fiber and ornamental crops, are vulnerable to FWB-causing *Fusarium* spp. Until more data are available, the cultivation, propagation, and trade of these species should be avoided in FWB-infested areas, as they may serve as unrecognized *Fusarium* spp. reservoirs that may inadvertently contribute to FWB dissemination into new areas. Moreover, *Heliconia* spp. have also been reported as hosts of the important banana bacterial pathogen *Ralstonia solanacearum* and banana bunchy top virus, which cause Moko and bunchy top disease, respectively (Blomme et al., 2017; Hamim et al., 2017). Therefore, we underscore the need for improved quarantine regulations, which is currently particularly relevant for the expansion of TR4 threatens the sustainability of global banana production.

## ACKNOWLEDGEMENTS

The authors would like to thank Vitropic (https://www.vitropic.fr) for providing Grand Naine plants used in this study. We thank the WUR-Unifarm personnel for their support during trials and for greenhouse facility maintenance.

## DECLARATIONS

EMP was supported by a NUFFIC PhD scholarship, grant number EPS 2016-02. GHJK and HJGM were supported by the Dutch Dioraphte Foundation. The funders had no role in study design, data collection and analysis, decision to publish, or preparation of the manuscript.

---

**Figure S1.**
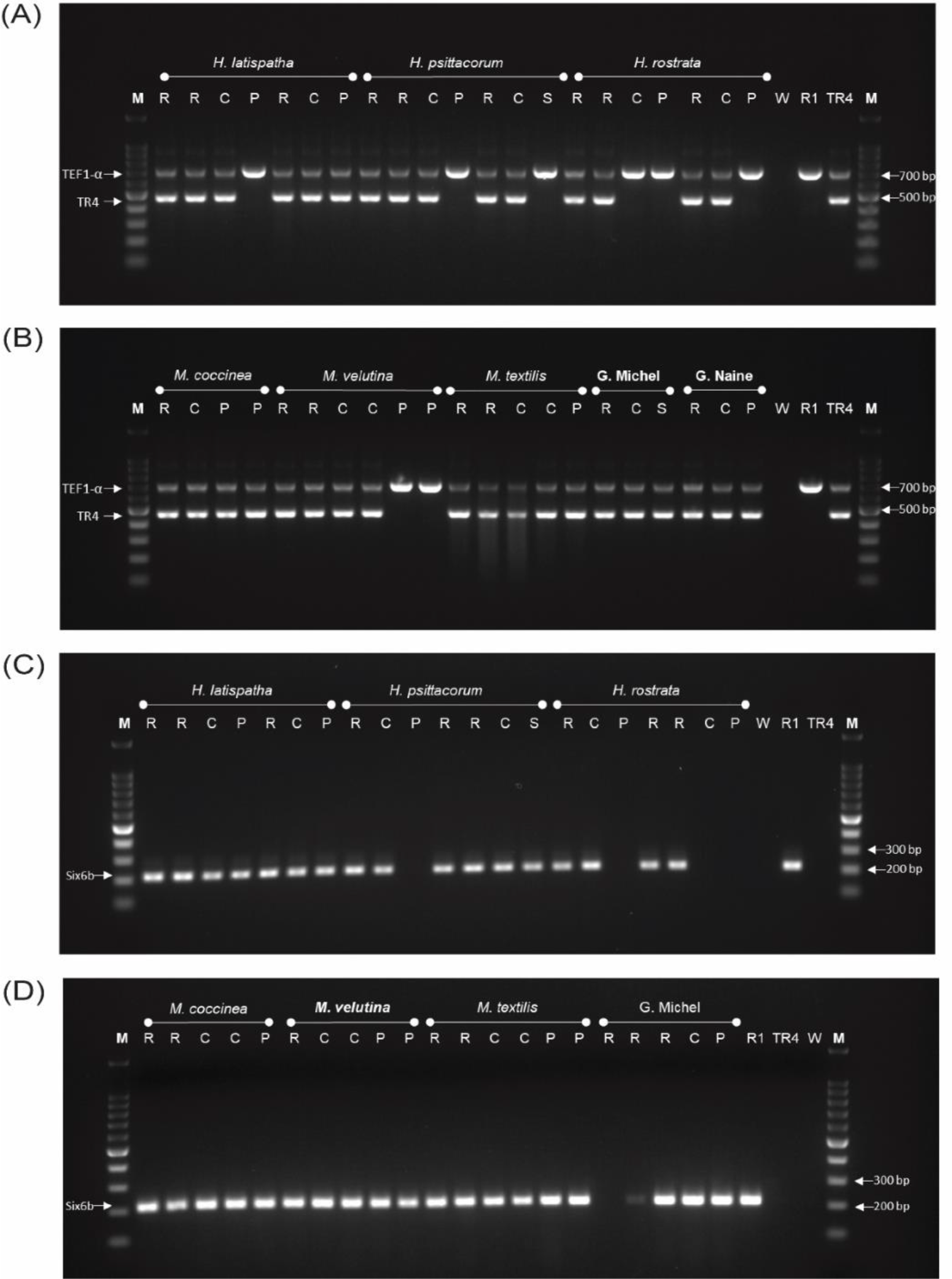
PCR identification of re-isolates obtained from different *Heliconia* and *Musa* species. (A) and (B) Agarose gel (1.5%) showing PCR amplicons following the molecular diagnosis of Dita et al., (2010), with the internal TEF1α amplicon and the TR4 diagnostic amplicon, corresponding with re-isolates from plants inoculated with TR4. (C) and (D) Agarose gel (1.5%) showing the results of PCR identification, according to the molecular diagnosis of Carvalhais et al., (2019), of the re-isolates obtained from plants inoculated with Race 1. Samples obtained from roots (R), corm (C), pseudostem (P). Controls comprise *Fusarium odoratissimum* strain II5 (TR4), *F. phialophorum* strain CR1.1A (R1) and water (W).

